# Prediction and functional interpretation of inter-chromosomal genome architecture from DNA sequence with TwinC

**DOI:** 10.1101/2024.09.16.613355

**Authors:** Anupama Jha, Borislav Hristov, Xiao Wang, Sheng Wang, William J. Greenleaf, Anshul Kundaje, Erez Lieberman Aiden, Alessandro Bertero, William Stafford Noble

## Abstract

Three-dimensional nuclear DNA architecture comprises well-studied intra-chromosomal (*cis*) folding and less characterized inter-chromosomal (*trans*) interfaces. Current predictive models of 3D genome folding can effectively infer pairwise *cis*-chromatin interactions from the primary DNA sequence but generally ignore *trans* contacts. There is an unmet need for robust models of *trans*-genome organization that provide insights into their underlying principles and functional relevance. We present TwinC, an interpretable convolutional neural network model that reliably predicts *trans* contacts measurable through proximity ligation-dependent (*in situ* and intact Hi-C) and independent (DNA SPRITE) genome-wide chromatin conformation assays.. TwinC uses a paired sequence design from replicate Hi-C experiments to learn single base pair relevance in *trans* interactions across two stretches of DNA. The method achieves high predictive accuracy (AUROC=0.80) on a cross-chromosomal test set from *in situ* and intact Hi-C experiments in heart tissue. Furthermore, we train TwinC using *in situ* Hi-C data from the widely used GM12878 cell line and validate its performance with orthogonal DNA SPRITE assay in the same cell type. Mechanistically, the neural network learns the importance of compartments, chromatin accessibility, clustered transcription factor binding and G-quadruplexes in forming *trans* contacts. In summary, TwinC models and interprets *trans* genome architecture, shedding light on this poorly understood aspect of gene regulation.

## 1 Introduction

The genetic information encoded in nuclear DNA is organized into a finely regulated 3D structure that controls tissue- and disease-specific gene expression [1, 2, 3]. *Cis* folding is well understood and comprises hierarchical structures such as DNA loops, topologically associating domains (TADs), and active/inactive (A/B) compartments. *Trans* interfaces are less widely studied. Non-holocentric chromosomes in higher eukaryotes occupy distinct domains in the interphase nucleus, referred to as “chromosome territories” [4]. Chromosome territories limit the frequency of *trans* interactions compared to the “spaghetti” DNA fiber model [5]. However, these territories do not serve as strict barriers. Certain loci can bypass these spatial restrictions, allowing interactions within different nuclear compartments, such as mRNA and tRNA factories, polycomb domains, the nucleolus, and nuclear speckles [6]. These interactions often involve specific loci crucial for gene regulation, particularly in enhancer hubs [7], transcription factories [8, 9, 10], and splicing factories [11].

Advances in chromosome conformation capture methods, e.g., Hi-C [12], micro-C [13], ChIA-PET [14], GAM [15], and SPRITE [16], enabled sequencing-based, high-throughput assays of chromatin interactions in many cell types and states. Although these technologies measure both *cis* and *trans* DNA contacts, analytical methods have focused primarily on the patterns and functions of *cis* genome folding. Notably, robust experimental measurement of *trans* contacts is challenging: these contacts typically contribute to only 20-30% of all measured pairwise DNA interactions and are spread across a much larger inter-chromosomal space [17] (Fig. 1A). Thus, *trans* contacts data is relatively sparse, and leveraging Hi-C data from biological replicates is crucial for identifying robust *trans* contacts in the presence of inherent sparsity.

**Figure 1:**
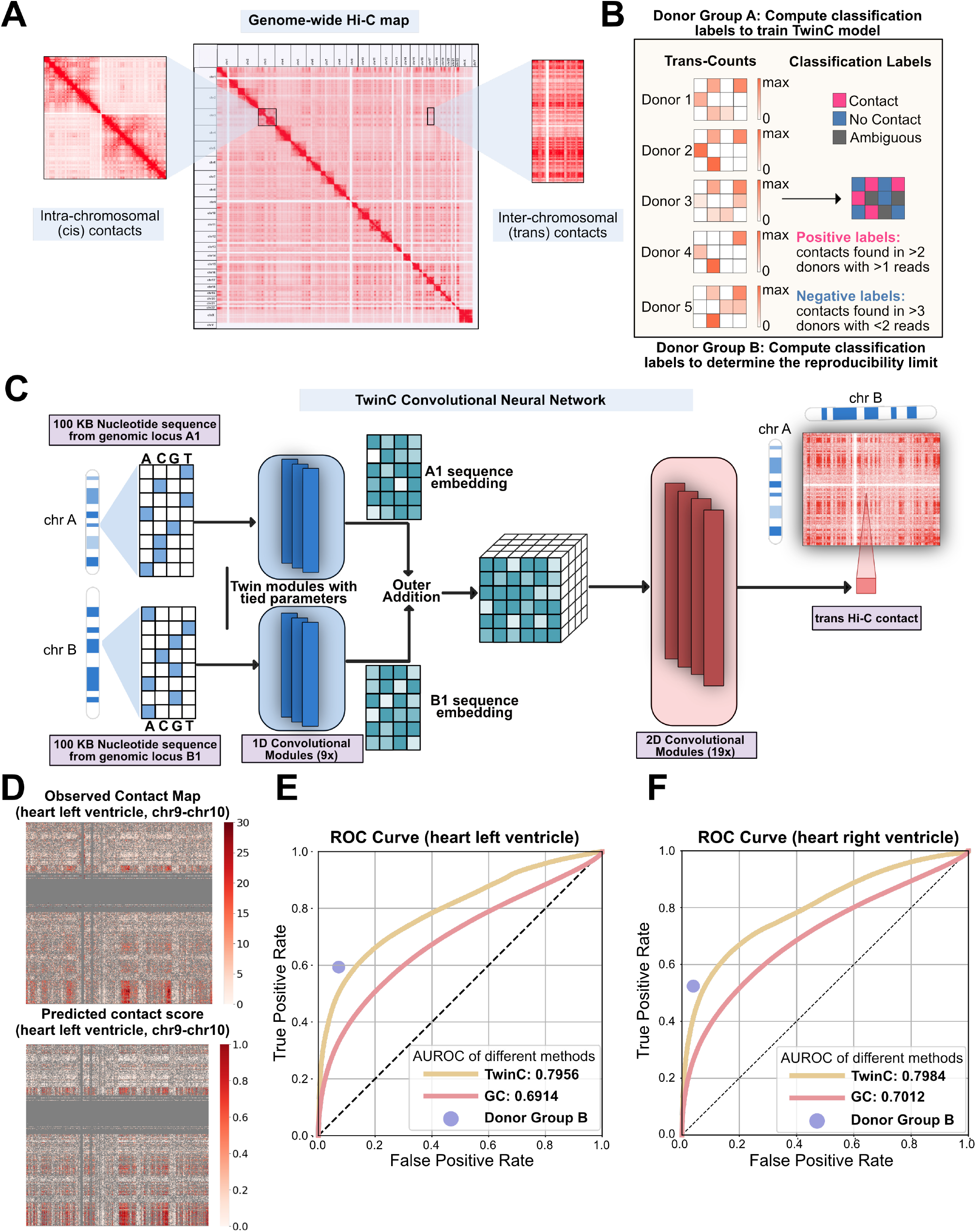
TwinC predicts reproducible *trans* contacts from nucleotide sequences in the human heart. **A**, A genome-wide Hi-C contact map with a focus on *cis* and *trans* contacts. **B**, The pipeline to extract labels using Hi-C contact maps from multiple donors. We divide the donors into two groups, donor group A and donor group B. Donor group A provides classification labels to train the TwinC model. Donor group B determines the upper limit of reproducibility between two donor groups by checking the agreement with labels from donor group A. **C**, Architecture of the TwinC model. **D–E**, ROC curves showing performance of TwinC (yellow curve) along with reproducibility upper limit (purple dot) and GC content baseline (red curve) in heart left ventricle (**D**) and right ventricle (**E**). **F**, Observed contact map (top) and predicted contact score from TwinC (bottom) from test set chromosomes chr9 and chr10 in heart left ventricle.

Machine learning methods can be leveraged to predict 3D genome folding and infer its underlying principles, including sequence and locus specificity. A first class of models, exemplified by Akita [18], DeepC [19], and Orca [20], learn to predict DNA contacts from DNA sequence alone and are evaluated on held-out chromosomes in the same tissue or cell type. Such models can be used to study the impact of sequence variations on DNA contacts within a tissue. A second class of models, including Epiphany [21] and C.Origami [22], learn to infer DNA contacts by combining DNA sequence with other epigenetic measurements like ATAC-seq, DNase-seq, or transcription factor (TF) ChIP-seq. These models can predict contacts on held-out chromosomes and in new tissue types, subject to the availability of the appropriate epigenetic data.

Both classes of models, however, are currently mostly restricted to predicting only *cis* contacts. Generally, these models take as input a single DNA sequence and produce as output the corresponding prediction of the contact frequencies for all pairs of genomic bins of a defined size within the given DNA sequence. This design only allows for the prediction of contacts between proximal pairs of DNA loci. Akita, for example, can only predict contacts at a resolution of 2048 basepair (bp) within 1 mbp of one another. This restriction is reasonable for *cis* interactions because chromatin’s polymer properties result in most observed *cis* contacts occurring within a 1 mbp window. However, extending such a model to capture all *cis* interactions requires a much larger receptive field, i.e., providing chromosome length sequences as input to the model.

Furthermore, generalizing this single-sequence setup to predict *trans* interactions is nontrivial. Currently, Orca is the only model that can be adapted to predict *trans* contacts: this is achieved by concatenating two 128 mbp sequences from different chromosomes as input to the model, employing a very large receptive field, and leveraging the model’s multi-resolution capabilities to zoom into the center of the input for predictions at 512 kbp, 256 kbp, and 128 kbp resolutions. This approach makes generating predictions for entire chromosome pairs at resolutions finer than 1024 kbp a complex and computationally expensive task.

To address these challenges, we present TwinC, a convolutional neural network (CNN) model specifically designed to predict *trans* DNA-DNA contacts. TwinC takes as input two DNA sequences, making the prediction of *trans* contacts straightforward. Furthermore, motivated by the extreme sparsity of typical *trans* contact maps, where even a single read-pair may be statistically significant if observed repeatedly, TwinC uses replicate Hi-C measurements in a given tissue type to determine whether a pair of DNA loci is repeatedly observed to interact (Fig. 1B). Accordingly, TwinC frames *trans* contact prediction as a classification task with “contact” versus “no contact” classes, in contrast to the regression setup of existing models. The TwinC code and trained model are freely available under an Apache license at https://github.com/Noble-Lab/twinc.

## 2 Results

### 2.1 TwinC, a twin convolutional neural network to predict *trans* genome folding

TwinC is a CNN model that predicts contact between two *trans* genomic loci from nucleotide sequences (Fig. 1C, Supplementary Figure 1). The model takes two 100 kbp nucleotide sequences as input and treats the task of predicting *trans* contacts as a classification task. Hence, TwinC outputs a “contact score,” a scalar value between 0 and 1, where 0 indicates the absence of contact, and 1 indicates the presence of contact between the input sequences. The high sparsity of *trans* contacts motivates the switch from the standard, regression-like framework to a classification setting. In the classification setting, we leverage multiple biological replicates to find reproducible contacts shared across replicates and use them as robust positive labels. Similarly, we leverage agreement between biological replicates regarding the absence of contacts to obtain reliable negative labels. Given two input sequences, the TwinC model first employs a 1D convolutional encoder to generate one sequence embedding for each given sequence. We chose 128 × 5 for the embedding dimension because it provides a slight performance benefit (Supplementary Figure 2). This encoder can capture local sequence motifs and their interactions, which are often relevant for genomic interactions. The two embeddings from the encoder are combined using outer addition. Outer addition between two *N* -dimensional tensors produces an (*N* + 1)-dimensional tensor where each element is the sum of one element from each input tensor, with broadcasting over all combinations of indices. The resulting tensor is passed to a 2D convolutional decoder, which integrates spatial interaction patterns between the two input embeddings and subsequently produces the contact score. By independently encoding two input sequences, TwinC can model long-range genomic interactions across arbitrary distances while maintaining a compact parameter footprint. For example, with 6.5 M parameters, TwinC is 3.5-fold smaller than the inter-chromosomal part of the Orca model with 22.5 M parameters. Further architectural details of the TwinC model are described in the Methods section.

### 2.2 TwinC accurately predicts reproducible *trans* contacts from nucleotide sequences in the human heart

We trained TwinC on Hi-C data from the left and right ventricles of the human heart, available from the ENCODE consortium. For each ventricle, we downloaded Hi-C experiments from ten human donors (Supplementary Table 1) and divided the donors into two groups, A and B. Robust positive and negative classification labels were extracted from each group (Fig. 1B). Group A labels were used to train TwinC, and group B labels were used to generate the upper bound of predictive performance. To estimate this upper bound, true positive and false positive rates were computed using labels from group A as ground truth and group B as predictions. These values represent the reproducibility between the two donor groups.

TwinC’s training procedure aims to minimize the binary cross-entropy loss function between the predictions and the targets. To avoid overfitting, we used five-fold validation, where in each fold, we divided the human genome with 19/2/2 chromosomes into the train/validation/test sets.

Our evaluation suggests that TwinC accurately predicts whether a pair of *trans* DNA loci participates in a robust contact. For predictions on test set chromosomes, TwinC achieves an AUROC of 0.7956 and 0.7984 in the left and right ventricles of the human heart, respectively (Fig. 1D-E). We compare TwinC to two competing methods. The first method, GC, is a simple baseline where we use the GC content of the input sequence pair as the contact predictor. The second method, Donor Group B, is a control that represents an upper bound on the predictive performance. TwinC substantially outperforms the GC baseline on the held-out set in the heart, left and right ventricle (Fig. 1D-E). Furthermore, TwinC’s performance is close to the upper bound in both tissues. In the heart left ventricle, TwinC achieves a true positive rate (TPR) of 0.4925, only 0.0959 below the TPR of the upper bound, 0.5884, at the same false positive rate (FPR) of 0.0662 (Fig. 1D). Similarly, in the heart right ventricle, at the FPR of 0.0482, TwinC’s TPR of 0.4430 is only 0.0691 below the upper bound TPR of 0.5121 (Fig. 1E). For a random classifier, the ROC curve lies on the line y = x. Hence, at a false positive rate of 0.0482, a random classifier is expected to achieve a true positive rate of 0.0482. Visually, TwinC captures a variety of patterns observed in the experimental Hi-C data (Fig. 1F). Furthermore, we evaluate the TwinC model using *in situ* Hi-C data in the heart left ventricle (Supplementary Table 2). Supplementary Figure 3 shows that the TwinC model, which is trained on intact Hi-C, achieves an AUROC of 0.8291 when using *in situ* Hi-C as labels compared to an AUROC of 0.7620 achieved by the GC baseline, highlighting the model’s ability to generalize across orthogonal chromatin conformation datasets that differ in their preservation of nuclear architecture.

To quantify potential biases in label generation resulting from donor selection, we randomly assigned five donors from the heart left ventricle Hi-C experiments to the label group and the remaining five to the prediction group, repeating this process four times. Subsequently, we trained four TwinC models, one for each donor group used for label assignment. Supplementary Figure 4 shows that the AUROC on the test set is consistent across four different label groups. This result demonstrates that our model’s performance is consistent regardless of which five donors were used in the label group. Finally, despite the propensity of smaller chromosomes to cluster together (Supplementary Figure 5A and 5B) [12], we found no bias in TwinC’s performance with respect to chromosome length (Supplementary Figure 5C and 5D). These results showcase TwinC’s ability to accurately predict robust *trans* contacts present in the ventricles of the human heart.

Notably, our training procedure is very efficient due to the small size of the TwinC model. We can train TwinC at 100 kbp resolution in 18 hours using one NVIDIA L40 40 GigaByte (GB) GPU. This training time represents a 26-fold reduction from Orca, where the third stage of the Orca model, which generates *trans* predictions at 128 kbp finest resolution, was trained for 20 days using a server with four NVIDIA Tesla V100 32 GB GPUs.

### 2.3 TwinC’s trans-contact predictions are validated with DNA SPRITE

For orthogonal validation of *trans*-contact predictions made by TwinC, we leveraged *in situ* Hi-C and DNA SPRITE experiments in the GM12878 cell line, a widely used cell line with several available 3D structure measurements (Supplementary Table 3) [23, 16]. *In situ* Hi-C uses the relative frequency of DNA-DNA proximity ligation events for genome-wide detection of pairwise interactions in the nucleus [24]. In contrast, DNA SPRITE uses split-pool recognition of interactions by tag extension for genome-wide detection of higher-order interactions within the nucleus without relying on proximity ligation [16]. However, the DNA SPRITE data can be converted from higher-order interaction clusters to a 2D contact map, and we used a previously generated 2D map [16] for our analysis. First, we trained TwinC using *in situ* Hi-C data in GM12878 using five-fold cross-validation with the same train/validation/test splits as the heart ventricles data (Fig.2A). Subsequently, we evaluated the TwinC model’s predictions on the test set chromosomes using DNA SPRITE data as labels. Fig.2B shows that at 1 Mb resolution (the resolution used by [16] for comparison between *in situ* Hi-C and DNA SPRITE) using DNA SPRITE data as labels, the agreement between the two experimental technologies is high (AUROC: 0.9611), and TwinC’s predictions are close to this upper limit (AUROC: 0.9336). This result shows that TwinC’s predictions generalize across experimental techniques for quantifying nuclear architecture based on fundamentally distinct principles.

### 2.4 TwinC’s accuracy is comparable to Orca’s on the task of *trans* contact prediction

To our knowledge, Orca is the only existing machine learning model capable of predicting *trans* contacts from DNA sequences. Accordingly, we set out to directly compare the two models. To enable a fair comparison, we implemented an alternate version of TwinC that mimics some of Orca’s design choices. First, we designed a regression version of TwinC because Orca frames the *trans* contact prediction task as a regression problem. Second, we modified TwinC to predict *trans* contacts at 128 kbp resolution, corresponding to the finest resolution at which the Orca model predicts *trans* contacts. Third, we trained the TwinC regression model on the H1 embryonic stem cells (H1ESC) cell line for which the trained Orca model is available, and we held out the identical chromosomes as the Orca test set, chromosomes 9 and 10. We generated 100 samples of 256 mbp multichromosomal sequences from test chromosomes 9 and 10 using Orca’s pipeline, and we repeated this sampling process ten times.

Comparing Orca’s and TwinC’s predictions to the held-out Hi-C data, we observe comparable test set performance between the two methods. Specifically, following Orca’s evaluation, we computed the Spearman correlation coefficient between normalized contact scores and predictions from the Orca and TwinC regression models for the H1ESC cell line (Fig. 3). When the observed contact data was normalized using Orca’s pipeline, both the Orca and TwinC models displayed similar correlation across ten groups of 100 samples each (Fig. 3A TwinC’s R > Orca’s R, in 6/10 samples and TwinC’s R < Orca’s R, in 4/10 samples) with good correspondence between normalized contact score and predictions from both models (Fig. 3B). Alternatively, when the observed contact data was normalized using TwinC’s pipeline, TwinC displays marginally better correlation than Orca (Fig. 3C, TwinC’s R > Orca’s R, in 10/10 samples, p-value=0.128 using Wilcoxon rank-sum test, Fig. 3D). These results suggest that TwinC performs comparably with the Orca model despite a simpler design and substantially faster training time.

**Figure 2:**
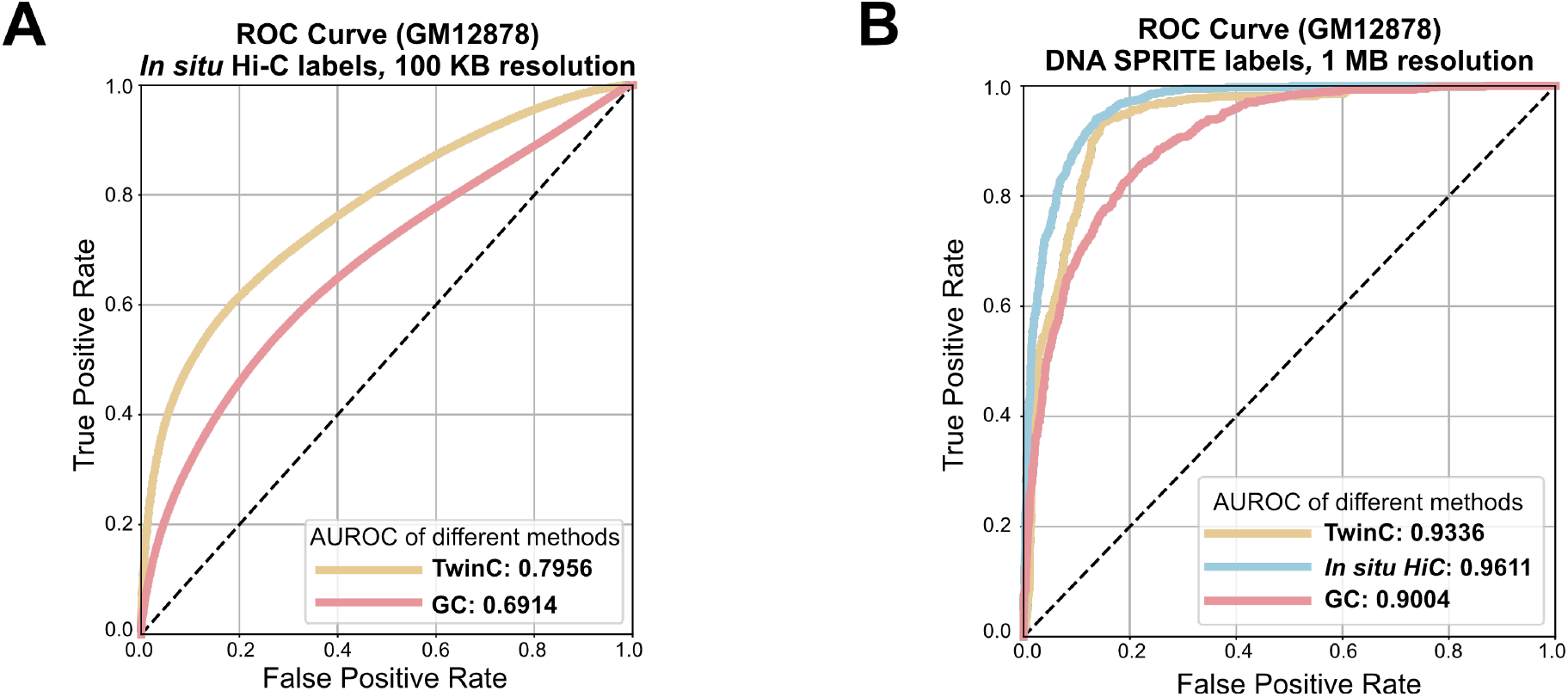
TwinC’s trans-contact predictions are validated with DNA SPRITE. **A**, ROC curves showing performance of TwinC (yellow curve) along with GC content baseline (red curve) on *in situ* Hi-C data in GM12878 cell line. **B**, ROC curve showing performance of TwinC (yellow curve) along with in situ Hi-C (blue curve) and GC baseline (red curve) when using DNA SPRITE data as ground truth labels in GM12878 cell line at 1 MB resolution.

**Figure 3:**
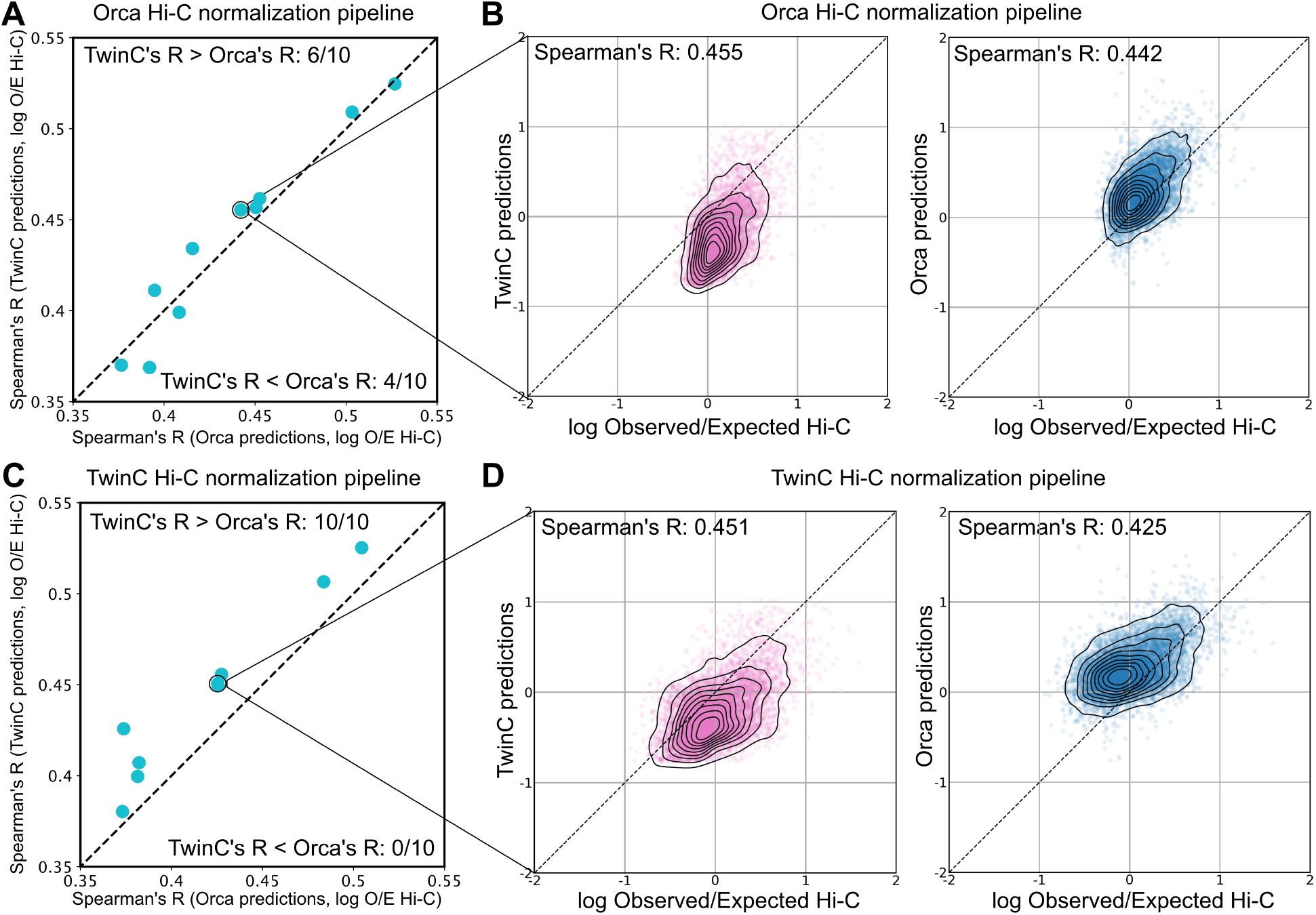
TwinC and Orca models perform at par on the task of *trans* contact prediction. **A**, Scatterplot showing Spearman’s R between Orca predictions and Hi-C data on the x-axis and Spearman’s R between TwinC predictions and Hi-C data on the y-axis. **B**, The left scatterplot shows Hi-C data on the x-axis and Orca predictions on the y-axis. The right scatterplot shows Hi-C data on the x-axis and TwinC predictions on the y-axis. The Hi-C data for **A-B** has been normalized using the Orca pipeline. **C-D** are similar to **A-B**, except the Hi-C normalization has been performed using the TwinC pipeline.

### 2.5 TwinC learns the importance of chromatin accessibility in the formation of *trans* contacts

A key goal in studying chromatin 3D architecture is to characterize the relationship between local chromatin structure and long-range chromatin contacts. We hypothesized that local chromatin accessibility is implicated in the co-localization of *trans* loci to facilitate regulatory processes like transcription in RNA factories [25]. Therefore, we designed a set of experiments to use TwinC to assess the impact of accessible DNA sequences on predicted *trans* contacts. The procedure uses ATAC-seq experiments to identify open chromatin regions, inserts or deletes the corresponding DNA sequences, and then uses TwinC to predict the resulting changes in *trans* contacts. Specifically, the first experiment consists of four steps. First, we downloaded ATAC-seq experiments in heart left and right ventricles from ENCODE and identified all peak regions therein. Second, we used TwinC to predict contact scores for all pairs of 100 kbp genomic loci between one test set chromosome pair, chr9 and chr10. Third, we ranked these locus pairs by the predicted contact score and designated two subsets, the top 10% and the next 30% of locus pairs (Fig. 4A). Fourth, for each top-10% locus pair, we moved all of its ATAC-seq peak sequences into the corresponding positions in a randomly selected locus pair from the next-30% set (Fig. 4B, illustration). We then compared the predicted TwinC contact score for these locus pairs before and after the substitution.

**Figure 4:**
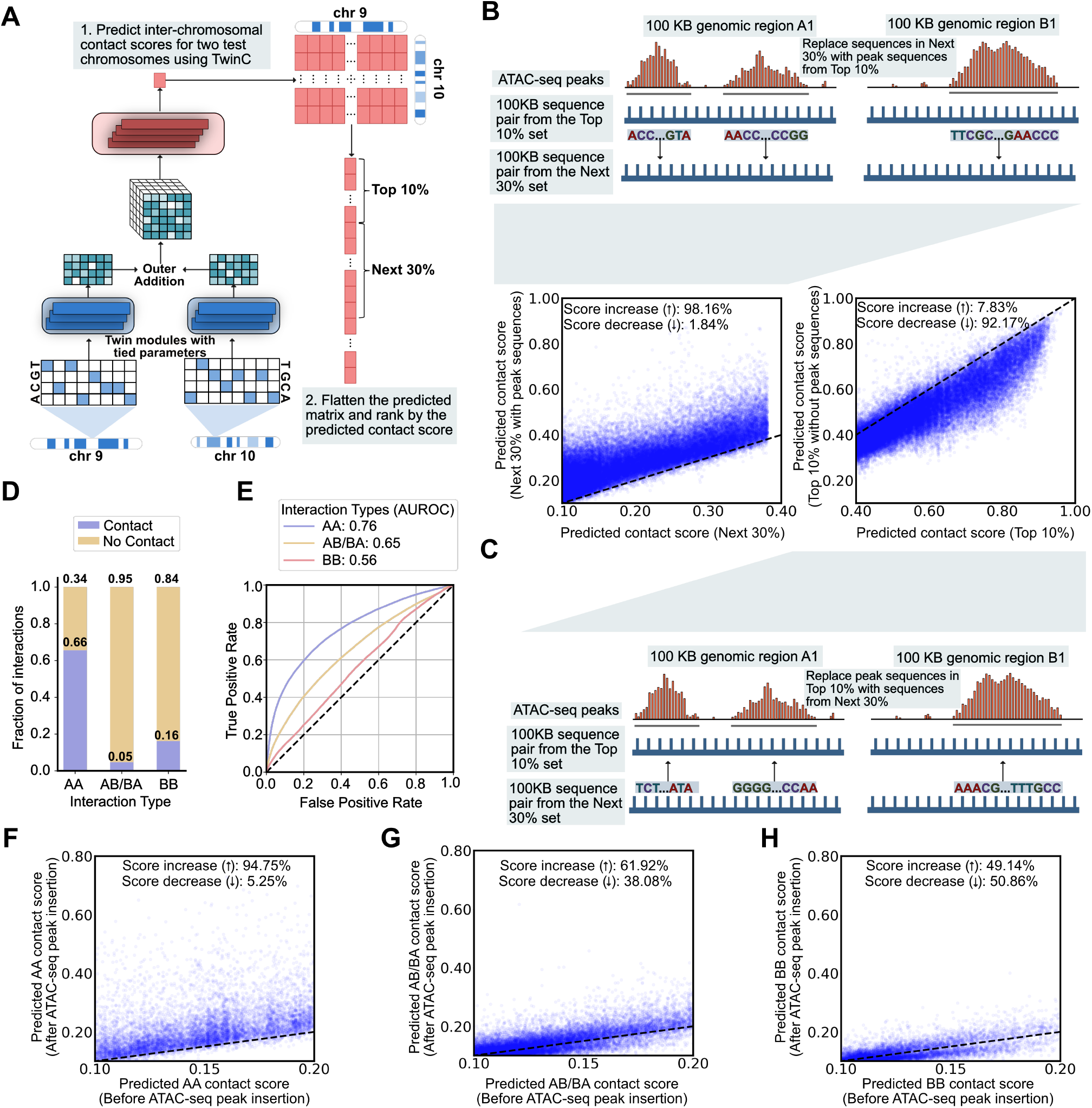
TwinC learns the importance of chromatin accessibility in the formation of *trans* contacts. **A**, Pipeline for generating a list of locus pairs from the test set chromosomes (chr9 and chr10), ranked by their predicted contact score. In the ranked list locus pairs, we designate two sets, top-10% and next-30%, which contain 10% and 30% of the locus pairs, respectively. **B**, Top illustration shows the experiment where DNA sequences corresponding to all ATAC-seq peaks in a random locus pair from the top-10% set are inserted into a random locus pair in the next-30% set. The bottom-left scatterplot shows the predicted contact score for the next-30% set before inserting peak sequences (x-axis) and after inserting the peak sequences (y-axis). **C**, The bottom illustration shows the reverse experiment, where DNA sequences from a random locus pair in the next-30% set are inserted in positions corresponding to all ATAC-seq peaks in a random locus pair from the top-10% set. The top-right scatterplot shows the predicted contact score for the Top 10% with peak sequences intact (x-axis) and without the peak sequences (y-axis). **D**, The stacked Bar plot showing the fraction of contacts versus no contact 100 kbp bin pairs in *trans* A-to-A (AA), A-to-B or B-to-A (AB/BA) and B-to-B (BB) bin pairs. **E**, ROC curve showing TwinC’s test set performance when predicting contacts based on compartment type: AA (Blue Curve), AB/BA (Yellow Curve) and BB (Red Curve). **F-H** The scatterplots show predicted contact score before insertion of ATAC-seq peaks from sequence pairs with higher predicted contact score (x-axis) and after insertion of these peaks (y-axis) for AA (**F**), AB/BA (**G**) and BB (**H**) compartment contacts.

This experiment shows that TwinC connects chromatin accessibility to *trans* contacts. When we insert ATAC-seq peak sequences into a randomly selected pair of loci, we observe an almost uniform increase in the predicted contact score: 98.16% scores increase, with a mean increase of 0.0855 in the predicted contact score (Fig. 4B, scatterplot). We repeated this experiment in the reverse direction, replacing ATAC-seq peaks in the top 10% set with corresponding sequences from the next 30% set (Fig. 4C, illustration). Again, we observed a marked shift in the predicted contact scores, but in the opposite direction (Fig. 4C, scatterplot). Together, these two results indicate that chromatin accessibility plays a role in TwinC’s *trans* contact predictions. As a corollary, this observation suggests that accessibility is a key feature of chromatin involved in *trans* contacts.

### 2.6 TwinC’s predictions differ systematically in A versus B compartments

Given the importance of A/B compartment structure in chromatin 3D architecture, we next investigated the distribution of reproducible contacts in A versus B compartments. First, we used POSSUMM [23] to assign A/B compartment labels based on the five heart left ventricle Hi-C experiments in our test set. For robustness, loci for which only three out of five experiments agreed on the compartment label were omitted. Using these compartment labels, we then segregated our *trans* contact labels into AA, AB/BA and BB compartment types (Fig. 4D). This analysis showed that *trans* contacts between A compartments are most frequent, followed by contacts between B compartments, with A-to-B contacts occurring least frequently.

Subsequently, we performed two experiments to investigate the difference in TwinC’s predictions by compartment type. First, we segregated positive and negative locus pairs into three categories (AA, AB/BA, BB) and separately calculated test set receiver operating characteristic (ROC) curves for each category (Fig. 4E). We observe a striking trend: AA compartment contacts are easiest to predict from DNA sequences (AUROC: 0.76), followed by AB/BA contacts (AUROC:0.65), with BB contacts being the most difficult to predict (AUROC: 0.56). This result suggests that TwinC successfully finds sequence-specific elements that mediate AA contacts but not BB contacts. This lack of sequence-specificity in BB contacts can be due to several reasons. For example, B compartments are enriched in repeat elements and mapping short reads from sequencing experiments to repeat regions is computationally infeasible, potentially resulting in our inability to distinguish contact regions enriched in repeat elements from non-contact regions enriched in repeat elements. Additionally, other layers of epigenomic regulation, such as histone modifications, could be involved. However, further experiments are required to test these hypotheses formally.

Second, we repeated the ATAC-seq peak insertion experiment described above but focused separately on AA, AB/BA and BB contacts. Once again, we observed a marked difference in TwinC’s behavior in these three settings. For AA contacts, inserting peak regions from sequences with higher contact scores to sequences with lower contact scores led to a uniform increase in their contact score (Fig. 4F), similar to what we observed previously in our aggregate analysis. However, when focusing on AB/BA and BB compartment contacts, this effect disappears; instead, the effect of the peak insertion leads to a nearly equal distribution of positive and negative changes in the predicted contact score (Fig. 4G-H). Together, these results indicate that contacts within the gene-rich A compartment depend more strongly upon accessible sequence elements than in the B compartment, which aligns with current models of these two compartments [26, 27].

### 2.7 TwinC finds transcription factors implicated in *trans* contacts

Aggregation of RNA polymerase II transcription initiation complexes in nuclear space is known to form transcription factories [28]. However, the role of individual TFs in these *trans* clusters is yet to be comprehensively characterized. Sequence-to-activity models like TwinC can contribute to this characterization task by quantifying the effect of TF motifs on *trans* contacts. To that end, we designed an experiment to quantify the importance of individual TF motifs for TwinC’s predictions (Fig 4A). We focused on 55,393 pairs of loci with the highest predicted contact scores from TwinC, and we used Integrated Gradients [29] to find per-base importance scores across all sequences within this set. We then examined the distribution of importance scores associated with a set of 1022 TF motifs derived from two sources, one downloaded from the JASPAR database and the second derived from the ChromBPNet model. Using the Mann-Whitney U test on the distribution of median importance scores associated with occurrences of each TF motif and a corresponding shuffled decoy motif, we identified 112 TFs that TwinC finds necessary for *trans* contacts (Fig. 5B, Supplementary Table 4).

**Figure 5:**
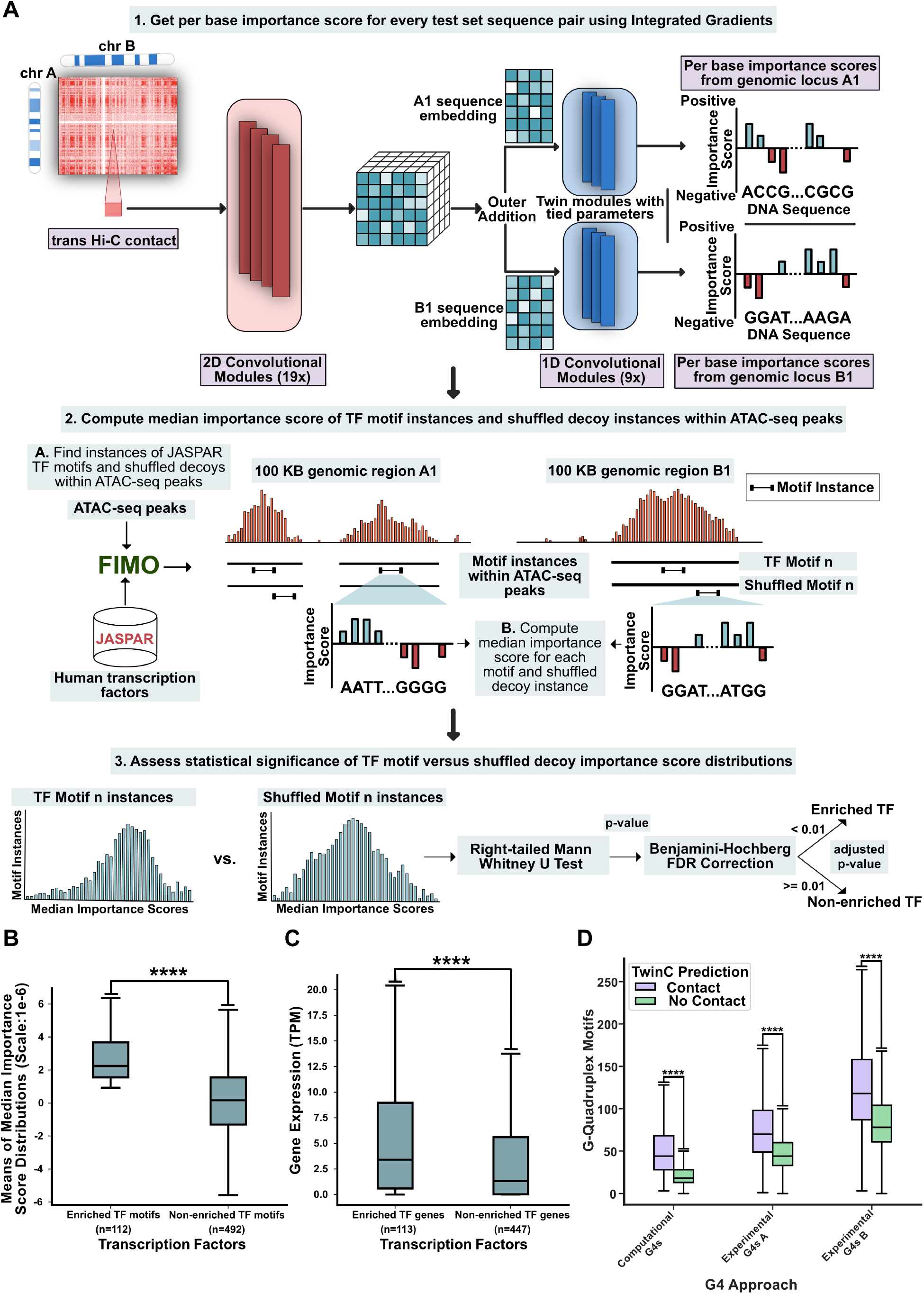
TwinC finds transcription factors enriched in *trans* contacts. **A**, Schematic showing the process of defining TFs enriched in *trans* contacts and non-enriched TFs using median importance score distributions of individual TF motifs and their shuffled decoys. **B**, Box plot showing means of median importance score distributions of enriched TF motifs (left, n=112) and non-enriched TF motifs (right, n=492). Per Base importance scores were generated using Integrated Gradients on the top 10% sequence pairs in the test set based on predicted contact score in the heart left ventricle. **C**, Box plot showing gene expression (TPM) of enriched TFs (left, n=113) and non-enriched TFs (right, n=447) in heart left ventricle. **D**, Paired boxplot showing the number of G-quadruplex motifs in predicted contact (contact score ≥ 0.5, in purple) and in predicted no contact (contact score *<* 0.5, in green) test set sequence pairs for three sources of G4 regions: computational G4 calls from quadparser and two G4-seq experiments. All p-values are from the right-tailed Mann-Whitney U test.

Our list of *trans* contact TFs includes several expected classes of motifs. For example, CTCF, with an adjusted p-value of 3.09 × 10^−309^, is a known regulator of *cis* genome architecture, essential for the formation of enhancer-promoter loops and enriched at TAD boundaries [30, 31, 32]. Previous studies provide specific instances of CTCF-mediated *trans* interactions, including Igf2/H19 and Wsb1/Nf1 loci [33] and in X chromosome inactivation in mammals [34, 35, 36, 37, 38]. Our results indicate a widespread prevalence of CTCF motifs in *trans* interactions. In addition, we observe the enrichment of several members of the ATF [39], TBX [40], RUNX [41], GATA [42], KLF/SP [43], ETS [44], IRF [45] and STAT [46] families of proteins, which are known to have key roles in cell types found in the heart, such as cardiomyocytes, endothelial cells, cardiac fibroblasts, macrophages and lymphocytes.

To investigate the properties shared by TFs enriched in *trans* contacts, we systematically assessed three properties of each TF: expression of the TF gene in heart ventricles, evolutionary conservation of the TF gene, and the presence of intrinsically disordered regions (IDRs) within the TF protein. We find the *trans* contact TFs are highly expressed (Fig. 5C) compared to a control set of TFs not enriched in *trans* contacts. However, there is no significant difference between the two groups in terms of evolutionary conservation and intrinsically disordered regions (Supplementary Figure 6A-D). Previous studies have shown that TFs must maintain a delicate balance between disorder, which can facilitate protein-protein interactions, and evolutionary selective pressure for stable structures [47, 48]. Our results suggest that TFs implicated in *trans* contacts may regulate their targets through higher expression but not necessarily differences in conservation or disorder in protein structure than non-enriched TFs.

Finally, we observed several G-rich motifs in the *trans* TF set, including some TFs, such as SP1, that bind to G-quadruplexes (G4s) [49]. G4s are secondary structures in guanine-rich nucleotide sequences implicated in modulating transcriptional activity [49]. We used two approaches to test the hypothesis that G4s are implicated in *trans* contacts. First, we used quadparser to search for G4-motifs in our test set [50]. In parallel, we downloaded G4 coordinates from the G4-seq protocol in the HEK293T cell line [51, 52, 53, 54]. Next, we separated the test set into contact (predicted contact score *>* 0.50) and no-contact (predicted contact score *<* 0.50) groups. Subsequently, we counted the number of G4 motifs overlapping with each sequence pair in both sets. We found significant enrichment of G4 motifs in the contact versus no-contact group using both approaches (Fig. 5D). In parallel, we found that the G4 motif counts derived from quadparser and G4-seq are reasonable predictors of *trans* contacts, with AUROC ranging from 0.638 to 0.655 (Supplementary Figure 7). While several studies have linked enhanced G4 formation with increased transcription activity [55, 49], our results suggest that proximity of G4-rich sequences in *trans* can be a potential mode of G4-mediated gene regulation.

## 3 Discussion

TwinC is a sequence-based CNN that learns to predict *trans* Hi-C contacts. Unlike existing approaches that predict all pairs of Hi-C contacts within a single genomic sequence, TwinC adopts a paired-sequence approach, predicting the contact frequency between two input sequences. In this regression setup, this approach results in a computationally efficient model with 3.5-fold fewer parameters, 26-fold faster training, and comparable predictive performance to the previous state-of-the-art Orca model. Due to extreme sparsity in *trans* contacts, TwinC moves away from the regression setup of predicting the frequency of Hi-C contacts to a classification setup where robust positive and negative labels are extracted from multiple replicate experiments. In this classification setup, TwinC achieves an AUROC of 0.7956 and 0.7984 in the heart left and right ventricles, respectively. Furthermore, TwinC’s TPR is close to the donor-based upper bound of predictive performance at the same FPR. Our results show the effectiveness of our paired-sequence approach in simplifying the architecture, training and inference of sequence-based 3D genome architecture models.

Using sequence substitution experiments, we showed that substituting ATAC-seq peak sequences from locus pairs with higher predicted contact scores to locus pairs with lower scores leads to an increase in contact score for the latter and vice versa. Furthermore, this effect is specific to A-to-A compartment contacts and disappears in B-to-B compartment contacts. Therefore, TwinC finds the sequence context of A compartment-specific accessible regions significant for *trans* contact formation and suggests that B-to-B compartment contacts may be regulated through other mechanisms. For example, previous studies have shown that B compartments overlap highly with Lamin-associated domains [23]. Therefore, the nuclear lamina may also contribute to their inter-chromosomal organization [56, 57].

TwinC finds 112 TF motifs implicated in *trans* contacts, using importance scores from Integrated Gradients combined with TF motifs from JASPAR and ChromBPNet. These *trans* contact TFs are highly expressed in the heart ventricles, and their motifs are highly predictive of *trans* contacts. Our list of TFs includes CTCF and several members of the ATF, TBX, RUNX, GATA, KLF/SP, ETS, IRF and STAT families of proteins, among others. Although previous studies have shown specific instances of CTCF-mediated *trans* interactions, our study suggests that the CTCF motif has a more widespread presence in *trans* contacts. Similarly, we have shown that the TFs TwinC finds to be enriched in *trans* contacts are highly expressed and functional in the heart, with their misregulation often implicated in cardiovascular diseases. However, the specific roles of these TFs in forming *trans* contacts have not yet been comprehensively characterized. Our study offers potential *trans* contact TF candidates for future studies looking into heart-specific or tissue-agnostic *trans* 3D genome architecture. TwinC also finds that G4s are enriched in *trans* contact. G4s are known to bind to transcription factors to enhance or repress transcription, and our study indicates that the proximity of G4-rich sequences in *trans* may be implicated in this regulation. However, more investigation is needed to establish the exact role of G4s, if any, in the formation of *trans* contacts.

In the current study, we applied TwinC to *trans* contacts, but this approach could also significantly simplify models for *cis* contacts. One future direction is to model *cis* and *trans* contacts together in one model using this approach. Such joint modeling can reveal the unified regulatory landscape of the 3D nuclear architecture. Furthermore, the sparsity of *trans* contacts due to the read coverage of Hi-C experiments is a major limitation for modeling *trans* interactions at finer resolution than 100 kbp. As higher coverage Hi-C experiments become more readily available, modeling interactions between chromosomes at finer resolution may reveal a fine-grained picture of *trans* nuclear regulation.

## 4 Methods

### 4.1 Hi-C data

In this work, we analyze intact Hi-C data from the left and right ventricles of the human heart (Supplementary Table 1). The data was downloaded from the ENCODE and contains ten samples each from the heart’s left and right ventricles from 20 different donors, with read coverage varying between 2.5 to 6 billion reads. For validation, we downloaded four *in situ* Hi-C samples in the heart left ventricle from the 4D Nucleome consortium (Supplementary Table 2). Finally, we analyze *in situ* Hi-C and DNA SPRITE experimental data in the GM12878 cell line (Supplementary Table 3).

The 4DN DCC processed the DNA SPRITE to generate pairwise contacts from the native multi-way contacts and stored them in the cool file format. Each Hi-C experiment produces a matrix (“contact map”), where each row and column corresponds to a fixed-size genomic locus called a “bin.” We refer to the size of these bins as the “resolution” of the Hi-C experiment. The value in the contact map at position (*i, j*) is the number of read pairs found as evidence of contact between bins *i* and *j*. The downloaded Hi-C datasets were already normalized using SCALE, a variant of the Knight-Ruiz (KR) matrix balancing algorithm.

### 4.2 Hi-C labels

*Trans* Hi-C contact maps suffer from sparsity along with biological and technical biases. To address these issues, we formulate the task of predicting *trans* contacts from nucleotide sequence as a binary classification task, where the positive class consists of pairs of *trans* genomic loci in contact, and the negative class consists of pairs of loci with no contact. To define the positive and negative sets of contacts, we randomly segregate ten replicate Hi-C matrices into two groups, with five experiments in the “labels” set and five in the “predictions” set. We subjected the “labels” set of matrices to a three-step pipeline to obtain class labels. The first step aims to eliminate bins with extremely large or small total counts because these counts can represent technical artifacts. To do so, we sum all five Hi-C experiments into a single matrix and then remove the top 0.1% and bottom 1% bins, ranked by total genome-wide *trans* contact counts. We remove these same bins from the five Hi-C experiments in the prediction group. The second step identifies robust contacts between pairs of *trans* bins to use as positive labels. Therefore, we identify contacts in the labels group observed with ≥ *i* count in ≥ *m* experiments and label these contacts as positives. The third step identifies robust no-contact locus pairs to use as negative labels. In this step, we iterate over all pairs of chromosomes and remove empty rows and columns from their pairwise contact map because these bins never interact with any bin from the second chromosome. This operation leaves us with *trans* loci with some potential for interaction. We then find contacts with ≦ *j* count in ≥ *n* experiments, and we label these contacts as negatives.

Thus, assigning positive and negative labels to a set of contact maps requires specifying five parameters: the bin size and the four thresholds listed above (*i, m, j* and *n*). To set these parameters, we systematically vary the five parameters to select their optimal values, generating an ROC curve for each parameter set by using the assigned positive/negative labels as ground truth and the contact counts from the prediction experiment as predictions. We selected the five parameters based on the corresponding AUROC values and selected a bin size of 100 kbp, and the four selected parameters are *i* = 2, *m* = 3, *j* = 1 and *n* = 4.

### 4.3 Chromosome-level training, validation and test sets

We perform five-fold cross-validation [58] and, in each fold, divide the 23 human chromosomes into training, validation, and test sets of 19, 2, and 2 chromosomes, respectively. We exclude the Y chromosome and mitochondrial DNA from our analysis. The validation set is used for hyperparameter selection for its corresponding fold, and each model is evaluated on its corresponding test set, averaging test set performance across folds. *Trans* contact maps were computed only between chromosomes within a set to avoid information leakage between the training, validation and test sets. The train, validation and test set sizes for different folds are listed in Supplementary Table 5 for the heart left ventricle and Supplementary Table 6 for the heart right ventricle samples. We store the nucleotide sequence from the GRCh38 version of the human genome as a one-hot-encoded memory map of size 3,088,286,401 × 4, where each row corresponds to a genomic position and each column corresponds to a nucleotide (*A* = 1, *C* = 2, *G* = 3, *T* = 4).

### 4.4 TwinC architecture

TwinC is a convolutional neural network that predicts *trans* contacts between two genomic loci from their nucleotide sequences. The input to the model is two one-hot-encoded, 100 kbp nucleotide sequences. Both input sequences pass through the same encoder, whose architecture is derived from the Akita [18] and Orca [20] models. The encoder consists of ten non-linear and ten linear 1D convolutional blocks, where each convolutional block consists of two 1D convolutional layers, each followed by a batch normalization layer (Supplementary Figure 1). Each linear convolutional block starts with a max-pooling layer, and a ReLU activation function follows each batch normalization layer in the non-linear convolutional block. The embeddings produced by the encoder are then concatenated and input to a decoder module modeled after Akita and Orca. The decoder consists of 19 non-linear and 19 linear 2D convolutional blocks, where each convolutional block consists of two 2D dilated convolutional layers, each followed by a batch normalization layer (Supplementary Figure 1). Like the encoder module, a ReLU activation function follows each batch normalization layer in the non-linear convolutional block. The output from the model is a contact score between 0 and 1. The TwinC model is implemented using the PyTorch framework.

### 4.5 TwinC training and evaluation

Formally, TwinC has two input matrices, *X*^1^ and *X*^2^, and one label vector, *Y*, where 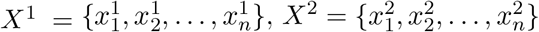, and 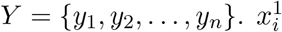 and 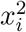 represent two one-hot-encoded sequence matrices, each of size 100,000 × 4, representing two nucleotide sequences from different chromosomes, and *y*_*i*_ is a scalar with a binary value of 0 for the absence of a Hi-C contact and 1 for the presence of a Hi-C contact between 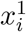 and 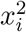, as determined using the procedure in Section 4.2. *Ŷ* represents the predictions from the TwinC model, where *Ŷ* = {*ŷ*_1_, *ŷ*_2_, …, *ŷ*_*n*_}. The TwinC neural network, represented by function *g*, produces each prediction 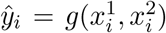 The training procedure optimizes a binary cross-entropy loss function:

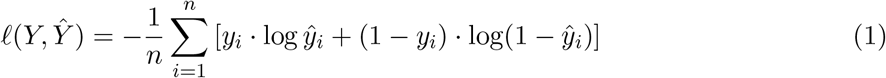

We use the PyTorch implementation of the Adam Optimizer for the training process. The initial learning rate of the optimizer is set to 1 × 10^−2^ with (*β*_1_, *β*_2_) = (0.9, 0.999) and no weight decay. The training data is input to the TwinC model in batches of size 16, and the model is trained for a maximum of 10 epochs. We evaluate the average precision on the validation data after every 200 batches of training data, and we save the current model if it achieves a better average precision than the previous best model. Because our training data are too large to fit into GPU memory, we customize the PyTorch DataGenerator object to generate training examples dynamically. This procedure involves uniformly sampling pairs of genomic loci from all training chromosomes, extracting the corresponding input sequences from a sequence memory map, and labels from a binary torch array. This optimization allows us to run TwinC on a single L40 GPU with 40 GB GPU memory.

### 4.6 Hyperparameter Selection

We estimated the optimal values for three hyperparameters using the performance on the validation set. First, we optimized the learning rate to achieve the best AUROC on the validation set and fastest convergence of training to be 1× 10^−2^ from the set of (1 × 10^−1^, 1× 10^−2^, 1× 10^−3^, 1× 10^−4^). Second, we tested five dimensions for the latent space (32 × 5, 64 × 5, 128 × 5, 256 × 5, 512 × 5) and observed that all five latent space dimensions resulted in similar predictive performance on the validation set (AUROC: 0.83-0.84, Supplementary Figure 2), with 128 × 5 latent dimensions resulting in a slight performance advantage. Therefore, we selected 128 × 5 as our latent space dimension. Third, we tested maximum pooling versus average pooling layers in our model and decided to use max pooling layers in our 1D and 2D convolutional modules based on the predictive performance on the validation set.

### 4.7 TwinC regression architecture

TwinC frames the *trans* contact prediction as a classification problem. In contrast, Orca solves a regression problem. Therefore, to enable a comparison between the two models, we implement a version of the TwinC model to solve the regression task. The TwinC regression model takes as input two sequences, each of length 640 kbp, and produces a 5 × 5 contact map output at 128 kbp resolution. TwinC regression follows the same encoder-decoder design as the classification model. The encoder consists of eight 1D convolutional block pairs, each consisting of a linear block followed by a non-linear block. Similarly, the decoder consists of 19 2D convolutional block pairs, each consisting of a linear block followed by a non-linear block.

### 4.8 TwinC regression training and evaluation

The TwinC regression model has two inputs, *R*^1^ and *R*^2^, and one target matrix, *T*, where 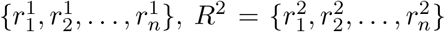, and *T* = {*t*_1_, *t*_2_, …, *t*_*n*_}. 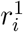 and 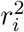 represent two one-hot-encoded sequence matrices, each of size 640,000 × 4, representing two nucleotide sequences from different chromosomes, and *t*_*i*_ is a 5 × 5 matrix representing the log normalized observed-over-expected *trans* contact map summarizing interactions between 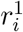 and 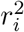 at 128 kbp resolution. 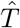 represents the predictions from the TwinC regression model, where 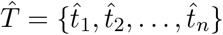. The TwinC regression neural network, represented by function *h*, produces each prediction 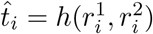 The training procedure minimizes a mean squared loss function:

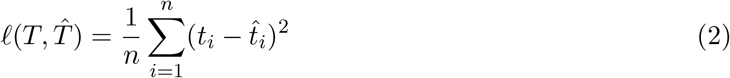

TwinC regression is implemented using the PyTorch library. We use the Adam optimizer for the training process with an initial learning rate of 9 × 10^−4^ with (*β*_1_, *β*_2_) = (0.9, 0.999) and no weight decay. Other training details of the TwinC regression model are identical to the classification model, described in Section 4.5.

### 4.9 Micro-C data processing

To enable comparison between the TwinC and Orca models, we download micro-C data in the H1ESC cell line, the same dataset that Orca was trained on, from the associated zenodo entry (https://zenodo.org/records/6234936). The downloaded data also included two trained Orca models and *trans* expected matrices at 32 kbp resolution from the H1ESC cell line. The micro-C dataset was already normalized using the Knight-Ruiz (KR) matrix balancing algorithm. Subsequently, to mitigate the impact of sparsity and technical artifacts, we filter bins from the training set in two steps. In the first step, we remove the top and bottom 0.01% bins ranked by total genome-wide *trans* contact counts from the training data. In the second step, we iterate over all pairs of chromosomes and remove empty rows and columns from their pairwise contact map because these bins never interact with any bin from the second chromosome. Then, we compute the observed-over-expected (o/e) *trans* contact ratios by dividing the observed contact frequency by the expected *trans* contact frequency over all chromosomes. Finally, we transform the o/e contact frequency using a natural log. We train and evaluate the TwinC regression model on the H1ESC micro-C dataset. Our train/validation/test split is identical to the one used by Orca: human chromosomes 9 and 10 are held out for testing, chromosomes 1 and 3 are used for validation, and the remaining 19 chromosomes are in the training set. We exclude the Y chromosome and mitochondrial DNA from our analysis.

### 4.10 Orca evaluation

Orca is a convolutional neural network that jointly predicts *cis* and *trans* contacts at multiple resolutions. Orca’s architecture comprises a multi-resolution hierarchical three-level encoder and a multi-level cascading decoder. The first level encoder takes 1 mbp sequences as input and predicts a 1000 × 1000 *cis* contact map at 1 kbp resolution. The second level encoder takes as input 32 mbp sequences and predicts a series of increasingly coarse-grained 250 × 250 *cis* contact map predictions at five resolutions from 4 kbp to 64 kbp. *Trans* interactions are allowed only at the third level, where the encoder takes an input sequence of 256 mbp and produces predictions at four resolutions from 128 kbp to 1024 kbp. To predict *trans* contacts, Orca randomly samples sequence segments between 64 mbp and 128 mbp from Orca’s test set chromosomes (chr9 and chr10). The samples are generated till a total length of 256 mbp has been sampled. The final sequence segment is truncated if the total length exceeds 256 mbp. The order of the sampled sequence segments is shuffled. Orca also takes loess-smoothed empirical expected scores at different resolutions as input. For *cis* contacts, the expected score at a genomic distance is the average normalized contact score across all chromosomes. The expected *trans* score is a scaler representing the genome-wide average normalized contact score. Therefore, Orca’s sampling process also makes a composite expected score matrix for four resolutions (128 kbp, 256 kbp, 512 kbp, 1024 kbp) composed of the *cis* expected matrix for each sampled sequence segment and a constant value that fills the expected matrix to represent interactions between sampled *trans* sequence segments. The 256 mbp multichromosomal input is then passed through the third-level Orca model, and *trans* positions in the predicted contact map are identified using the composite expected matrix. We follow the steps outlined in the Orca manuscript and the provided Python code to generate *trans* predictions. One hundred samples of 256 mbp multichromosomal input sequences and distance encoding matrices are generated using Orca’s code. Multiscale predictions for each input are produced by zooming in to the center of the 256 mbp sequence. The sampling process is repeated ten times, and we evaluate the Orca model at 128 kbp, its finest resolution for *trans* predictions.

### 4.11 Compartment calling

To identify A and B compartments, we ran the POSSUMM [23] tool on five donor Hi-C experiments at 100 kbp resolution. POSSUMM returns the first four eigenvectors for every chromosome based on PCA of the corresponding *cis* Hi-C contact map, using GC content to ensure that positive eigenvalues correspond to GC-rich regions of the genome. To avoid “compartments” that correspond to entire chromosome arms, we additionally use ATAC-seq data in left and right heart ventricles for validation. The ATAC-seq counts are averaged at 100 kbp resolution, and their correlation with the first eigenvector by POSSUMM is computed. We select the first eigenvector if it correlates well with ATAC-seq data (PCC > 0.30). Otherwise, the second eigenvector is selected. The first eigenvector passes the ATAC-seq validation in four out of five experiments. Once the eigenvectors from all five donors are generated, the compartment type for a given 100 kbp bin is decided by majority vote. For robustness, locus pairs resulting in a 3-to-2 vote are discarded.

### 4.12 Chromatin accessibility sequence substitution experiments

To assess the impact of local chromatin accessibility on trans contacts, we perform two experiments. The first experiment involves inserting ATAC-seq peak sequences into sequence pairs that are less likely to form contacts, and the second experiment involves removing ATAC-seq peak sequences from sequence pairs that are more likely to be in contact with the nucleus. For the first experiment, TwinC predictions on the test set chromosomes (chr9 and chr10) are first ranked by the predicted contact score. From this list, we identify two subsets: the top 10% and the next 30% of locus pairs. Next, we downloaded IDR-ranked peaks from 29 ATAC-seq experiments in the ENCODE portal (Supplementary Table 7), which contain 15 BED files from the heart’s left ventricle and 14 BED files from the right ventricle. The peaks are merged and sorted into a single BED file per tissue using bedtools. Subsequently, a random locus pair from the top 10% and the next 30% sets are selected. For each such pair, ATAC-seq peaks are intersected with the two 100 kbp regions corresponding to the locus pair from the top 10% set. Next, we insert sequences corresponding to such peaks from the locus pair from the top 10% set into identical positions in the locus pair from the next 30% set. Then, TwinC predicts the contact score of the modified sequence pairs from the next 30% set. For the second experiment, we perform the reverse sequence replacement operation: replacing the peak sequences in a locus pair from the top 10% set with identical length sequences from the same index position in a locus pair from the next 30% set. TwinC predictions are generated for the modified input sequence pairs from the top 10% set.

### 4.13 Finding enriched transcription factors using Integrated Gradients

We identify TFs implicated in *trans* contacts in five steps. First, we compile a set of 1022 TF motifs, which includes tissue-specific and tissue-agnostic motifs. This set consists of 821 human TF motifs from the JASPAR database (Supplementary Table 8) and 191 TF motifs generated by applying ChromBPNet [59] and TF-MoDisco [60] to heart ATAC-seq data (Supplementary Tables 9-12). The data comprises pseudobulk ATAC-seq datasets covering four cell populations in the human heart ventricles: cardiomyocytes, smooth muscle cells, fibroblasts and endothelial cells. Four ChromBPNet models predict chromatin accessibility from these four datasets using DNA sequences as input. Importance scores are generated for each cell population using the interpretation method DeepLIFT, and then TF-MoDisco extracts the top 38 to 67 TF motifs in each of the four cell types (Supplementary Tables 9-12). The JASPAR and ChromBPNet motifs are combined to produce a set of 1022 motifs for subsequent analysis. Second, we extract the top 10% (55,393) test set sequence pairs, ranked by their TwinC predicted contact score, and use FIMO [61] with a p-value threshold of 1e-5 to identify motif occurrences within accessible peaks present in these regions. Fifteen ATAC-seq experiments are merged to obtain the accessible peaks in the heart left ventricle (Supplementary Table 7). Similarly, peak coordinates from 14 ATAC-seq experiments are merged to extract accessible peaks in the heart right ventricle (Supplementary Table 7). Third, we generate a background set of motif occurrences by shuffling the columns of each motif and searching with FIMO against the same set of accessible regions. Fourth, we use Integrated Gradients to compute per-base importance scores for each motif occurrence in the foreground and background sets, assigning the median importance score to each occurrence. Fifth, we test the hypothesis that the median importance scores are higher among the real versus background motif occurrences using a right-tailed Mann-Whitney U test, including Benjamini-Hochberg correction with *α* = 0.01.

### 4.14 RNA-seq expression

We downloaded processed RNA-seq data in 435 heart left ventricle samples from the GTEx portal [62]. The table contains TPM counts for human genes processed with the GRCh38 gene annotations. Using the right-tailed Mann-Whitney U test, we evaluate the expression difference between TFs enriched and not enriched in *trans* contacts.

### 4.15 Intrinsically disordered regions

To find the intrinsically disordered regions (IDR), we download the amino acid sequences for all TFs from Uniprot (https://www.uniprot.org/proteomes/UP000005640). Subsequently, we use IUPRED3 [63] to predict per-base IDR scores and the number of globular domains for each TF. A per-base IDR score is also generated for every TF using ANCHOR2 [64]. We use the right-tailed Mann-Whitney U test to evaluate the difference in average IDR score and the number of globular domains between TFs enriched and not enriched in *trans* contacts.

### 4.16 G-quadruplexes

We adopt two approaches to find G-quadruplexes in the test set sequence pairs. First, we use Quadparser, a computational tool, to find the G4 consensus motif, *G*_3+_*N*_1−7_*G*_3+_*N*_1−7_*G*_3+_*N*_1−7_*G*_3+_, where G represents guanine, and N represents any nucleotide. We use the Quadparser implementation from the bioinformatics-cafe repository to carry out the motif search (https://github.com/dariober/bioinformatics-cafe/tree/master/fastaRegexFinder). Second, we obtain G4 coordinates in HEK293T cell lines in BED format from four experiments (GEO accession IDs: GSE63874, GSE110582, GSM3003539, GSM3003540). Next, we overlap the G4 coordinates obtained using Quadparser and G4-seq with the test set sequence pairs and count the number of G4 instances within each pair. Subsequently, we divide the test set sequence pairs into contact (predicted contact score *>* 0.5) and no-contact (predicted contact score ≦ 0.5) groups and compare G4 motif enrichment in the two groups using the right-tailed Mann-Whitney U test. Finally, we used the G4 motif counts to predict *trans* contacts. We computed AUROC using motif counts as the predicted score and the observed binary labels from the test set as the ground truth.

## Data availability

All data used in this study is publicly available, with accession numbers provided in the supplementary tables.

## Code availability

The TwinC code and trained model are available under an Apache license at https://github.com/Noble-Lab/twinc.

## Acknowledgments

The authors thank Salil Sanjay Deshpande for compiling the list of transcription factors enriched in the accessible regions across different cell populations in the heart ventricles.

## Author contributions

Conceptualization: A.J. and W.S.N., Data curation: A.J., W.S.N., E.L.A. and A.B., Formal analysis: A.J., W.S.N., A.B., X.W., and B.H., Funding acquisition: W.S.N. and A.J., Investigation: A.J., W.S.N., and A.B., Methodology: A.J. and W.S.N., Project administration: W.S.N, Resources: W.S.N., Software: A.J., Supervision: W.S.N., A.B., W.J.G, E.L.A, A.K., and S.W., Validation: A.J., A.B. and W.S.N., Visualization: A.J., Writing – original draft: A.J., W.S.N., and A.B. with input from all authors, Writing – review & editing: A.J., W.S.N., and A.B. with input from all authors.

