## Supplementary figures for "Prediction and functional interpretation of inter-chromosomal genome architecture from DNA sequence with TwinC"

### Supplemental information for predicting interchromosomal Hi-C contacts from DNA sequence with TwinC

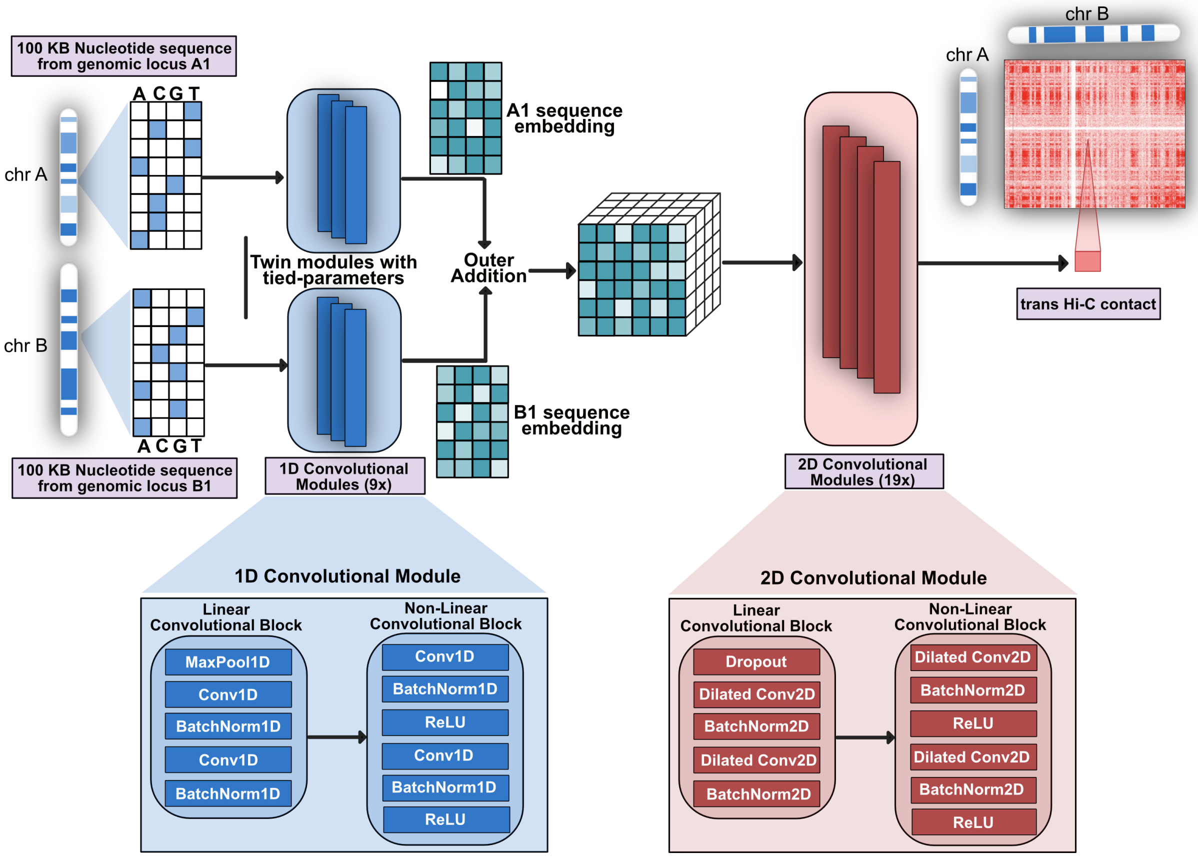


**Supplementary Figure** 1: **TwinC model architecture**. TwinC takes two 100 KB sequences that go through twin 1D convolutional encoder modules with tied parameters as input. Outer addition is performed on the two sequence embeddings from the encoder, and the output goes to the 2D convolutional decoder module to produce a predicted contact score between 0 and 1. The encoder and decoder layer designs derive from the previous 3D genome architecture models, Akita [2] and Orca [4].

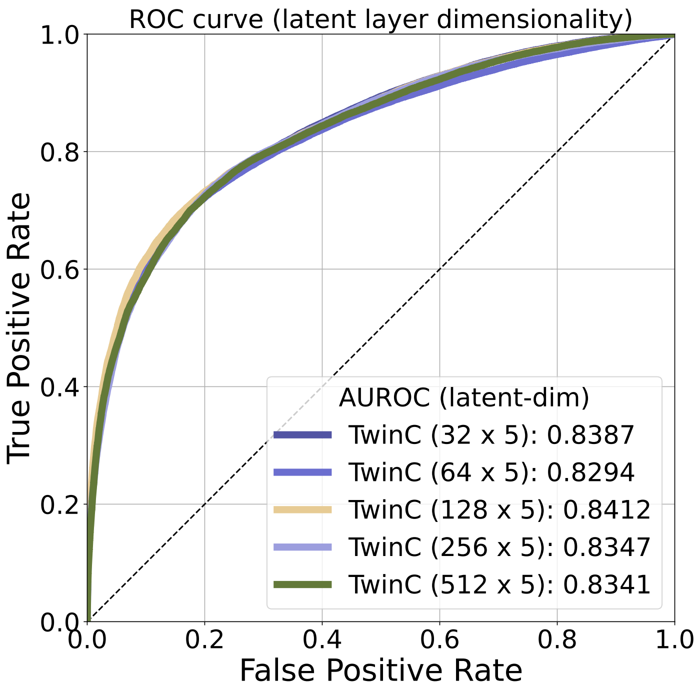


**Supplementary Figure** 2: **Testing different latent layer dimensions for TwinC.** ROC curve showing performance of TwinC using five different latent layer dimensions: $32 \times 5$, $64 \times 5$, $128 \times 5$, $256 \times 5$ and $512 \times 5$.

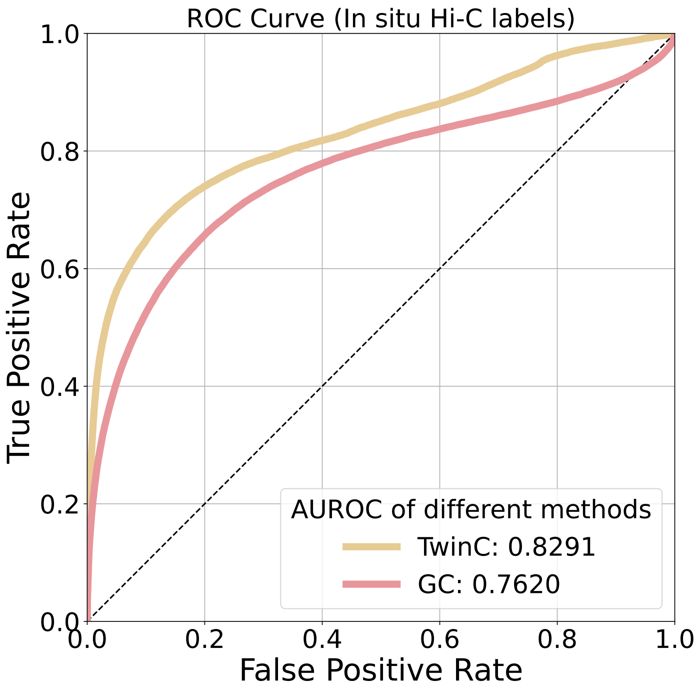


**Supplementary Figure 3: Evaluating TwinC model trained using intact Hi-C on *in situ* Hi-C data in heart left ventricle.** ROC curve showing performance of TwinC (yellow curve) along with GC baseline (red curve) when using *in situ* Hi-C data as ground truth labels at 100 KB resolution.

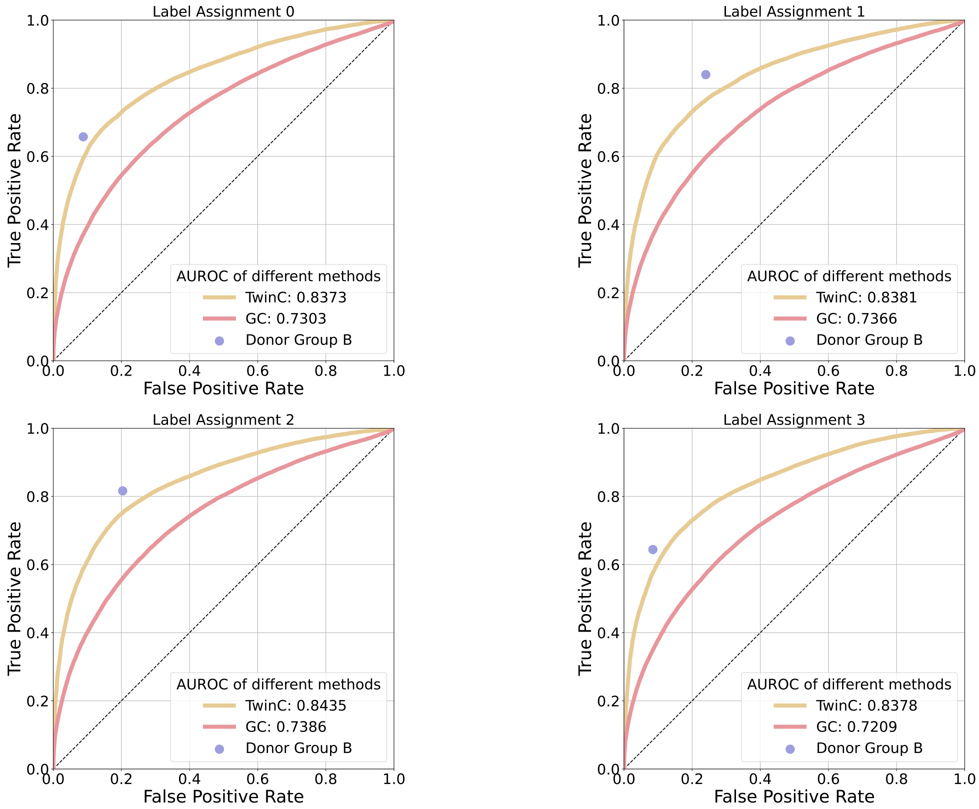


**Supplementary Figure** 4: **Testing label randomness introduced by donor selection.** Four ROC curves showing the performance of TwinC (yellow curve) along with the reproducibility upper limit (purple dot) and the GC content baseline (red curve) in the heart's left ventricle. Each ROC curve uses labels generated by a different subset of five donors (referred to as label assignments 0-3).

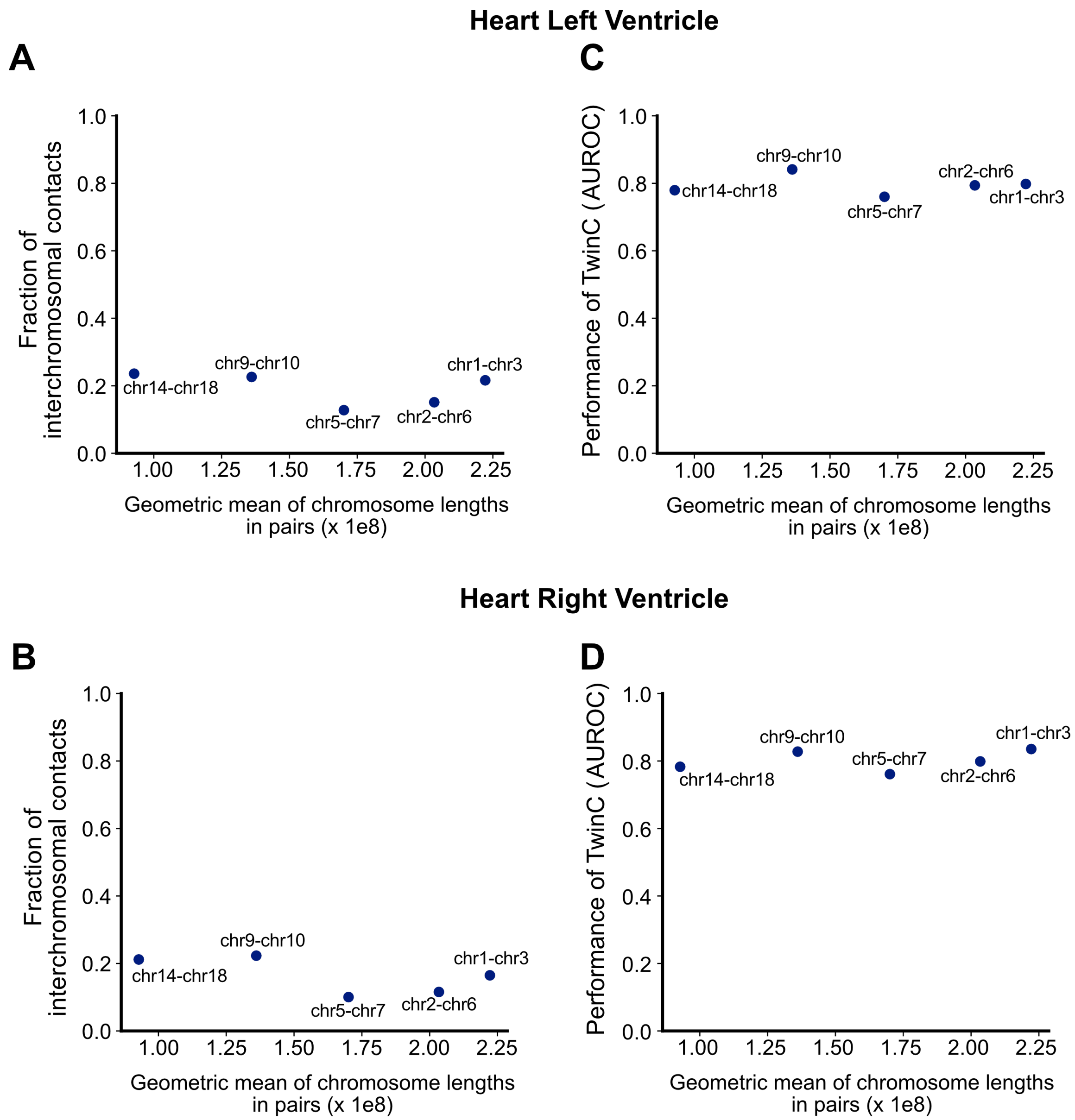


**Supplementary Figure** 5: **TwinC's performance based on chromosome length A-B,** Scatterplots showing the geometric mean of length of each chromosome in the test set chromosome pairs (x-axis) and the fraction of contacts in all 100 KB bin pairs in each chromosome pair (y-axis) in heart left (**A**) and right (**B**) ventricles. Each dot shows the names of the two chromosomes. **C-D,** Scatterplots showing the geometric mean of length of each chromosome in the test set chromosome pairs (x-axis) and AUROC score of TwinC on the corresponding test set chromosome pairs (y-axis) in heart left (**C**) and right (**D**) ventricles.

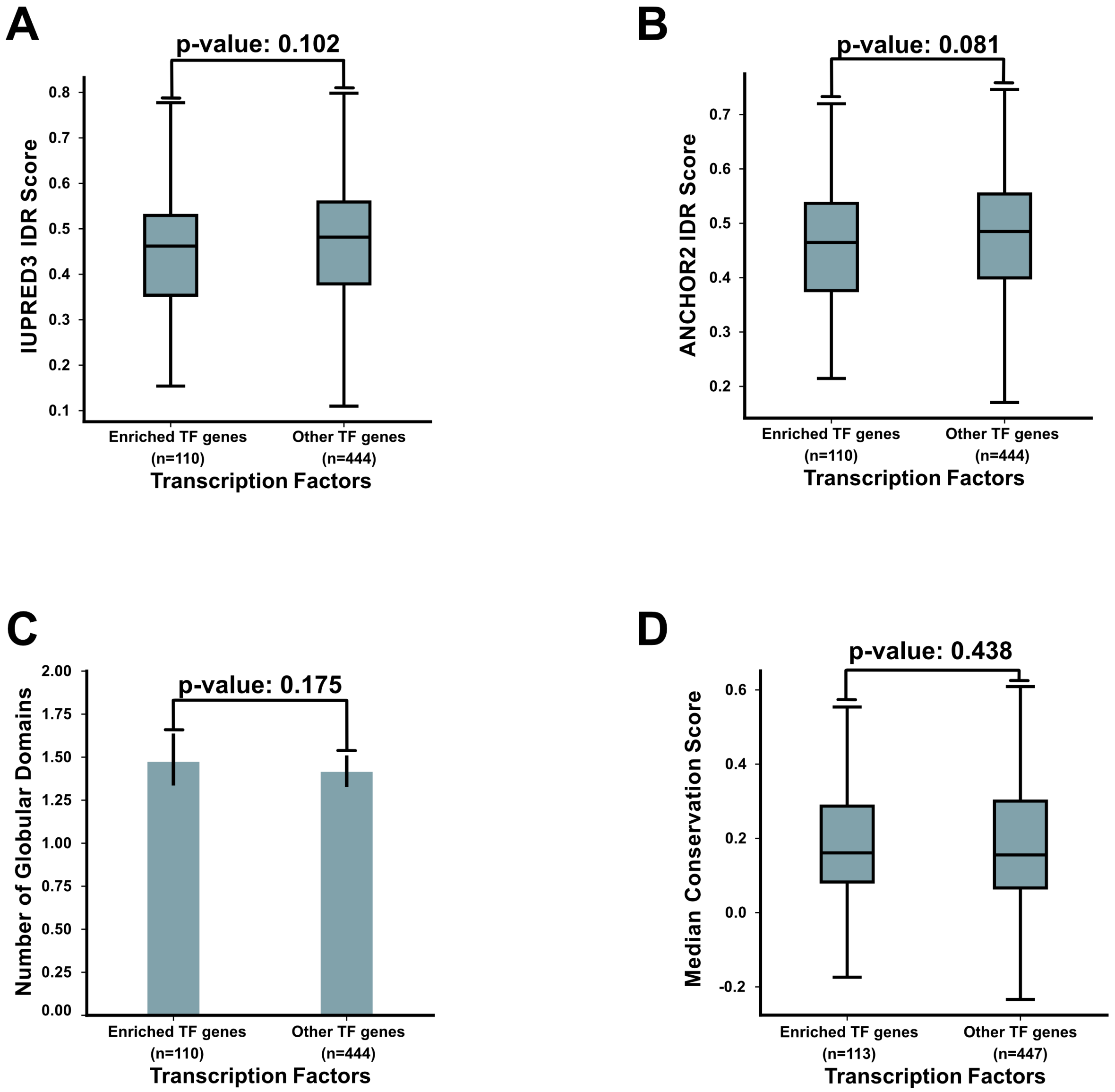


**Supplementary Figure** 6: **Disorder and conservation status of TwinC's interchromosomal contact TFs. A-B,** Box plot showing median intrinsically disordered regions (IDR) score from IUPRED3 [1] (**A**) and ANCHOR2 [3] (**B**) for TFs enriched in interchromosomal contacts (n=110) and other TFs (n=444). P-values in **A-B** from left-tailed Mann-Whitney U test. **C,** Bar plot showing the number of globular domains in TFs enriched in interchromosomal contacts (n=110) and other TFs (n=444). **D,** Box plot showing median PhyloP conservation score for enriched TFs (n=113) and other TFs (n=447). P-values in **C-D** from right-tailed Mann-Whitney U test.

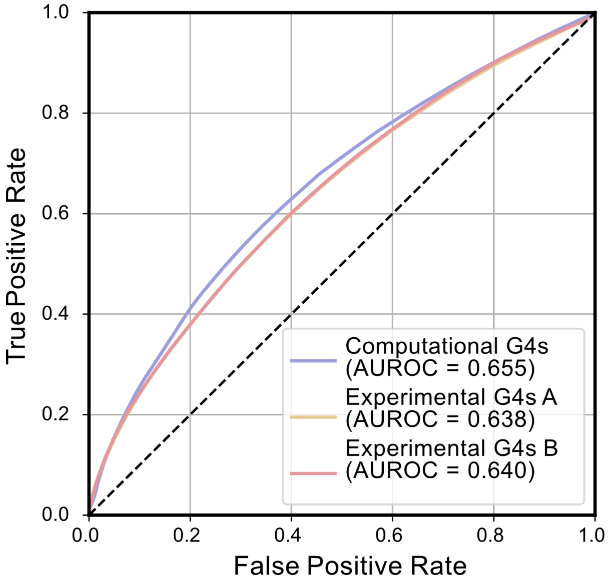


**Supplementary Figure 7: TwinC finds G-quadruplexes implicated in interchromosomal contacts.** ROC curve showing predictive performance when using the number of G4 motifs from three sources, computational G4 calls from quadparser and two G4-seq experiments as individual predictors of interchromosomal contacts on the test set.

#### References


[1] Gábor Erdős, Mátyás Pajkos, and Zsuzsanna Dosztányi. Iupred3: prediction of protein disorder enhanced with unambiguous experimental annotation and visualization of evolutionary conservation. *Nucleic acids research*, 49(W1):W297–W303, 2021.

[2] Geoff Fudenberg, David R Kelley, and Katherine S Pollard. Predicting 3d genome folding from dna sequence with akita. *Nature Methods*, 17(11):1111–1117, 2020.

[3] Bálint Mészáros, István Simon, and Zsuzsanna Dosztányi. Prediction of protein binding regions in disordered proteins. *PLoS computational biology*, 5(5):e1000376, 2009.

[4] J. Zhou. Sequence-based modeling of three-dimensional genome architecture from kilobase to chromosome scale. *Nature Genetics*, 54:725–734, 2022.
